# Efficient In Vitro Regeneration from Cotyledon Nodes and In Planta Genetic Transformation in Elite Peanut Cultivars

**DOI:** 10.64898/2026.02.11.705295

**Authors:** Charli Kaushal, Priyanka Rajput, Harini Gowrishankar, Mansi Parekh, Mahak Sachdev, HN Karthik, Liya Philip, Mukesh Jain, Subramanian Sankaranarayanan, Bhuvan Pathak

## Abstract

Peanut (*Arachis hypogaea* L.), a vital oilseed and food legume, is cultivated across the globe. Genetic improvement via conventional breeding faces limitations from narrow diversity and reproductive barriers, underscoring the need for tissue culture-based regeneration and transformation platforms. This study optimizes an efficient, reproducible in vitro regeneration protocol using cotyledonary node explants from three Indian elite cultivars: GG-20, GJG-9, and TAG-37A. Explants from aseptically germinated seedlings were cultured on Murashige and Skoog (MS) medium with varying cytokinins (e.g., BAP 0–4 mg/L) and auxins (e.g., NAA 0.1-0.9mg/L), yielding direct multiple shoot induction without callus, minimizing somaclonal variation. Optimal shoot proliferation occurred on full-strength MS + 2 mg/L BAP for GG-20/GJG-9 (88.9% efficiency) and 4 mg/L BAP for TAG-37A (∼89–100% efficiency); rooting peaked on half-MS + NAA (up to 88.9% in GG-20). Regenerated plants acclimatized successfully in greenhouse conditions. Additionally, a robust *in planta Agrobacterium tumefaciens* (EHA105, pGFPGUSPlus) transformation via plumular meristem pricking in GG-20 achieved 7.69% efficiency. Transgene integration was confirmed by GUS assay and PCR (GUS/hptII), with ∼64% soil establishment.

## Introduction

Peanut (*Arachis hypogaea L*.), an allotetraploid (AABB; 2n = 4x = 40) oilseed crop, holds major economic importance worldwide due to its high-quality edible oil. Cultivated on ∼28.9 million hectares across 100 countries, it produced 51.31 million tonnes in 2024 (Kaushal et al., 2025). In Western countries, peanuts are primarily consumed as confectionery and snacks, while in the Indian subcontinent, they serve as cooking oil and confectionery sources. Kernels contain 40–55% fat, 20–30% protein, 10–20% carbohydrates, plus essential nutrients like vitamins E and B-complex, and minerals (zinc, iron, calcium, magnesium, phosphorus, potassium) (Dean et al., 2009; Shasidhar et al., 2020). This nutrient profile positions peanut as a key player in combating malnutrition in Africa and Asia.

Although conventional breeding has played a major role in peanut varietal improvement, its pace remains insufficient to meet the growing demand for rapid cultivar replacement. Genomics-assisted breeding (GAB), particularly marker-assisted backcrossing (MABC), has enabled the efficient introgression of key traits such as foliar disease resistance, high oleic acid content, and nematode resistance in peanut (Chu et al., 2011; Varshney et al., 2014; Janila et al., 2016a; Kolekar et al., 2017; Bera et al., 2018; Shasidhar et al., 2020). Several molecular breeding lines developed through these approaches have demonstrated enhanced pod and haulm yield, with some released as cultivars and others under advanced stages of evaluation (Janila etl., 2016b; Shasidhar et al., 2020). However, the impact of these modern breeding strategies is often constrained by long generation cycles and the recalcitrant nature of cultivated peanuts.

Plant tissue culture offers a complementary strategy by enabling rapid *in-vitro* regeneration, clonal propagation, and providing a foundation for genetic transformation and genome-editing applications. Despite its potential, reproducible and genotype-responsive regeneration protocols for elite peanut cultivars remain limited. Indian peanuts are available in different varieties: Bold (Runner), Java (Spanish) and Red Natal types, varying in oil/protein content, seed size/ weight, and gene expression (Gawande et al., 2026). Gujarat, India’s leading peanut-producing state, widely cultivates varieties such as GG-20 and GJG-9, TAG-37A.

The GG-20, a Virginia bunch variety released in 1992, and developed from the cross GAUG-10 × R-33-1 is a high-yielding, semi-spreading peanut variety adaptable to semi-arid regions (Bharodia et al., 1997; Mahatma et al., 2020). It matures medium-duration, producing medium to large pods with two to three seeds and the pod yield of 1960 kg ha□^1^ and 50.7% oil content reported (Rathnakumar et al., 2013; Shasidhar et al., 2020). Its traits support studies in conventional agronomy and in vitro assays. GJG-9 (Gujarat Junagadh Groundnut 9) was included as a representative of drought-tolerant Spanish Bunch genotypes (Mahatma et al., 2020; Faldu et al., 2018). TAG-37A is a peanut variety from the cross TG25 × TG26, officially notified in 2004 for multiple Indian agro-climatic zones. It has a growth duration of 100–105 days, smooth pods with moderate seed size, and good drought tolerance. Suitable for kharif and rabi/summer cropping, it yields 2500–3000 kg/ha. Its short duration, drought resistance, and adaptability contribute to its popularity and status as a key check variety in comparative agronomic studies (https://www.barc.gov.in/group/66_s246.pdf). Its agronomic performance and seed quality are strongly influenced by genetic background, environmental adaptability, and agronomic practices, making the choice of cultivar critical for both field performance and in vitro responses in tissue culture studies (Mahatma et al., 2020). Cultivar choice critically influences field performance and tissue culture responses due to genetic/environmental/agronomic interactions.

Transferring genes into plants requires an effective *In-vitro* regeneration system. Various studies report on peanut micropropagation techniques, including mature cotyledons (Venkatachalam et al., 1999; Maina et al., 2010), nodals (Joboury et al., 2012), de-embryonated cotyledon (Radhakrishna et al., 2000; Pratap et al., 2018) and cotyledonary nodes (Verma et al., 2011; 2014; Limbua et al., 2019). Additionally, organogenesis (Pestana et al., 1999), somatic embryogenesis, and callus culture have been explored in peanut and other crops also (Iqbal et al., 2011; Mayavan et al. 2015). However, peanut cultivars exhibit significant variability in response to culture media, limiting the use of a single nutrient medium for optimal regeneration across different cultivars (Hoa et al., 2021). *In planta* transformation offers a somaclone-free alternative (Manickavasagam et al. 2015). Pioneered in *Arabidopsis* (Feldmann and Marks, 1987), this approach has been successfully applied to several recalcitrant crops, including chickpea and pigeonpea, with methodological modifications to enhance transformation efficiency (Chakrabarty et al., 2000; Ganguly et al., 2018). Rohini and Rao (2000) first reported *in-planta* transformation in peanut using the mature seed embryo axis, achieving a transformation efficiency of 3.3%. However, this method was not tested across different cultivars. In the study by Karthik et al. (2018), the cotyledon with an intact embryo was used as the explant for evaluating in planta transformation efficiency. Transformation was carried out using *Agrobacterium* strain EHA105 harboring pCAMBIA-based vectors, and stringent BASTA® selection was employed to effectively eliminate chimeric and non-transformed plants. Using this optimized approach, transformation efficiencies ranging from 31.3% in cv. CO6 to 38.6% in cv. TMV7 were achieved.

This study reports an efficient *in-vitro* regeneration system using cotyledonary node explants for elite peanut varieties (GG-20 GJG-9 and TAG-37A) to facilitate tissue culture-assisted varietal improvement. Further, we developed a robust and high-efficiency *in-planta* transformation protocol for peanut cv. GG-20, with the objective of achieving enhanced transformation efficiency, reliable plant recovery. Therefore, supporting functional genomics studies and biotechnological interventions aimed at improving oil quality and other agronomically important traits.

## Material and methods

### Plant Material

Studies were performed with peanut (*Arachis hypogaea* L.) cultivar GG-20, GJG-9, TAG-37A. GG-20 was provided by Junagarh Agricultural University, Gujarat, India. GJG-9 and, TAG-37A were provided by AlpGiri Seed Sciences Pvt Ltd, Gujarat, India. GG-20 (Virginia bunch, released 1992, high-yielding with 1960 kg/ha pods and 50.7% oil), GJG-9 (drought-tolerant Spanish bunch), and TAG-37A (short-duration, 100–105 days, 2500–3000 kg/ha yield, drought-resistant).

### Seed Sterilization and In Vitro Germination

Healthy peanut seeds (n=65 per variety) of three varieties-GG-20, GJG-9, TAG-37A (Fig1; Fig. 2A; Fig. 4A; Fig. 6A) were thoroughly rinsed under running tap water and soaked for 3-4 hrs. Following the initial rinse with sterile distilled water, the seed coats were aseptically removed, and the kernels were washed again with sterile distilled water for 2-3min. Kernels were surface sterilized with 70% (v/v) ethanol for 15-20 second and 0.1 % (w/v) HgCl_2_ solution for 2-3 min, and washed 4-5 times in sterile distilled water for 5 min each. Sterile seeds were dried in laminar air flow for 15-20 min and germinated on the half or full-strength basal MS (Murashige and Skoog 1962) medium containing 3% (w/v) sucrose and 0.8% clarigel (w/v), pH of 5.8 (Fig. 1; Fig. 2A; Fig. 4A; Fig. 6A; Table. 1). After 2 days of the incubation in the dark condition, seeds started to germinate (Fig. 2B; Fig. 4B; Fig. 6B). We selected the 50 most robustly germinated plantlets ones for subsequent experiments to maintain the genetic uniformity and stability of the experimental material.

**Figure 1.**
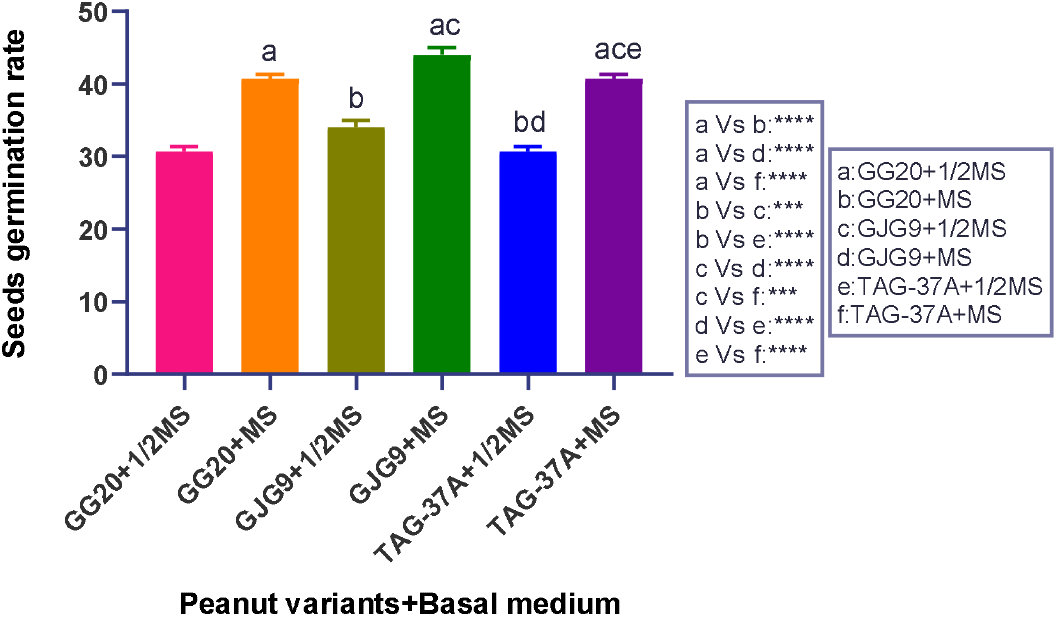
Effects of Basal medium (MS) on seed germination rate of three different variants on peanut variates (*Arachis hypogea*); GG-20, GJG-9, TAG-37A. MS: Murashige and Skoog medium; Bars with different letters indicates significant differences (*p* ≤ 0.05), as analysed by one-way ANOVA and Tukey’s multiple range test

**Figure 2.**
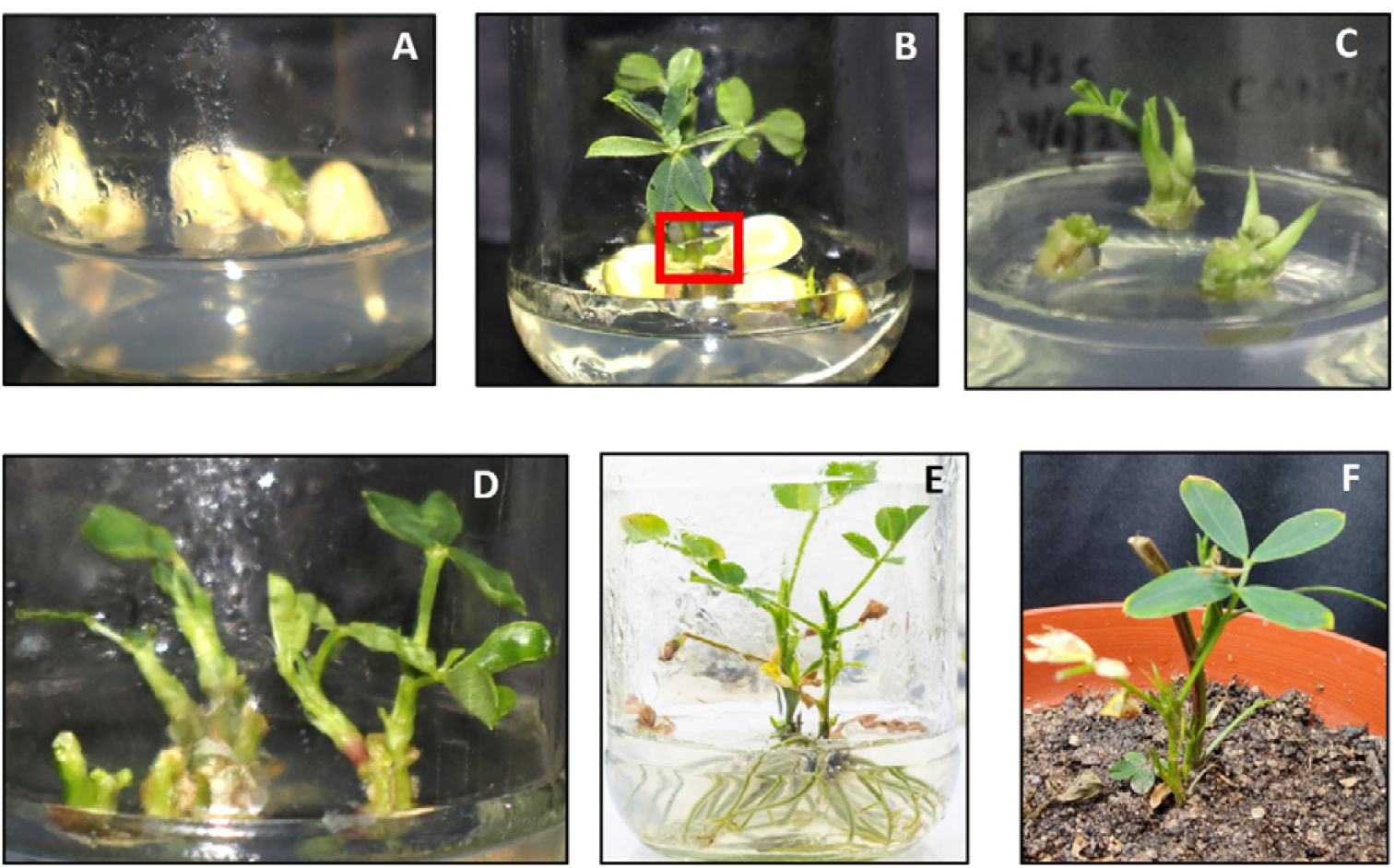
Tissue culture and hardening of *GG20* peanut cultivar; **(A)** Sterilized *A. hypogaea GG20* seed on MS medium; germination starts; **(B)** cotyledon node used for regeneration after seed germination; (Red mark represents that this portion excised by cutting and used as a (CNs) cotyledon node); **(C)** Cotyledon nodes regenerate on MS+ 2mg/L BAP medium; **(D)** Multiple regenerated shoots were ready for rooting; (E) Regenerated shoots were used for rooting on MS+ 0.9 mg/L NAA medium; **(F)** Rooted plants transferred in pots for the hardening purpose.

**Table 1.** Optimization of seed germination rate of different peanut variants (*Arachis hypogea*); GG20, JJ20, TAG-37A.

| S. No | MS media | Name of variant | Efficiency |
| --- | --- | --- | --- |
| 1. | ½ Basal Media | GG-20 | 60% |
| 2. | Full Basal Media | GG-20 | 80% |
| 3. | ½ Basal Media | GJG-9 | 70% |
| 4. | Full Basal Media | GJG-9 | 90% |
| 5. | ½ Basal Media | TAG-37A | 60% |
| 6. | Full Basal Media | TAG-37A | 80% |

### Explant Preparation

After the 2 days of the incubation the seedlings were transferred under 24□°C, 16 h photoperiod and 130–150 µmol m^−2^ s^−1^ florescent light conditions. The explant, cotyledon nodes (CNs) were excised used from 8 days old *in vitro* germinated plants (Fig. 2C; Fig. 4C; Fig. 6C) (Hsieh et al., 2017). Different concentration of cultures of shoot regeneration and multiplication, and root induction were maintained at 25±2ºC at the cool white, the fluorescent light intensity of 27 μmol/m2/s with 12 h daylight. *In vitro* growth in all experiments was evaluated at week 3-4 of culture.

### Optimization of shoot induction and elongation, and root induction media

CNs explants (8 days old) were cultured on MS medium supplemented with various concentrations of benzylminopurine (BAP) from 0-4 mg/L to induce shoot in all variants (Fig. 2C; Fig. 3A; Fig. 4C; Fig. 5A; Fig. 6C; Fig. 7A) (Table. 2,4,6). After the 8 days of shoot were further transferred on same media for multiplication (Fig. 2D; Fig. 4D; Fig. 6D). After the 10-12 days of the shoot’s multiplication the single shoot from the shoot clusters transferred on MS medium containing naphthaleneacetic acid (NAA) from 0-0.9mg/L or in combination with 0-3mg/L BAP for rooting (Fig. 2E; Fig. 3B; Fig. 4E; Fig. 5B; Fig. 6E; Fig. 7B) (Table. 3,5,7). The experiment was conducted with three replicates with nine CNs. After one month, the acclimatation rooted plants were transferred to pots containing mixture of sand: vermiculite: perlite 2:2:1 respectively with some gypsum and small amount of commercial rhizobium suspension culture in the green house (Fig. 2F; Fig. 4F; Fig. 6F).

**Figure 3.**
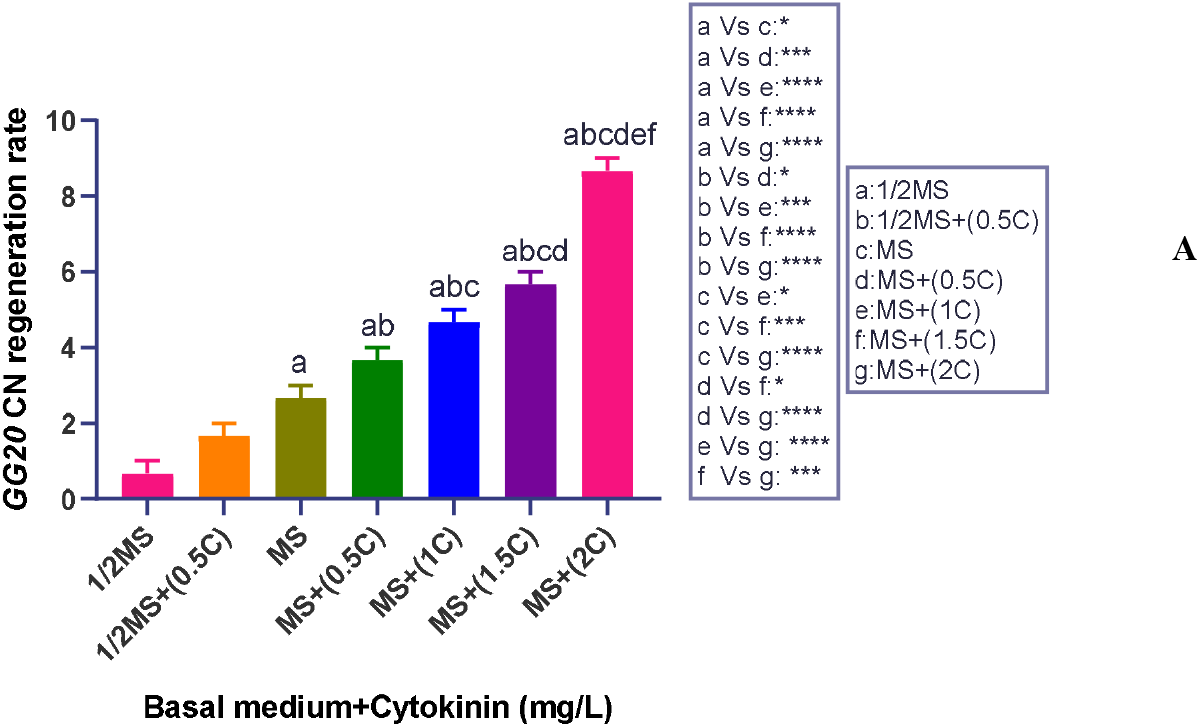
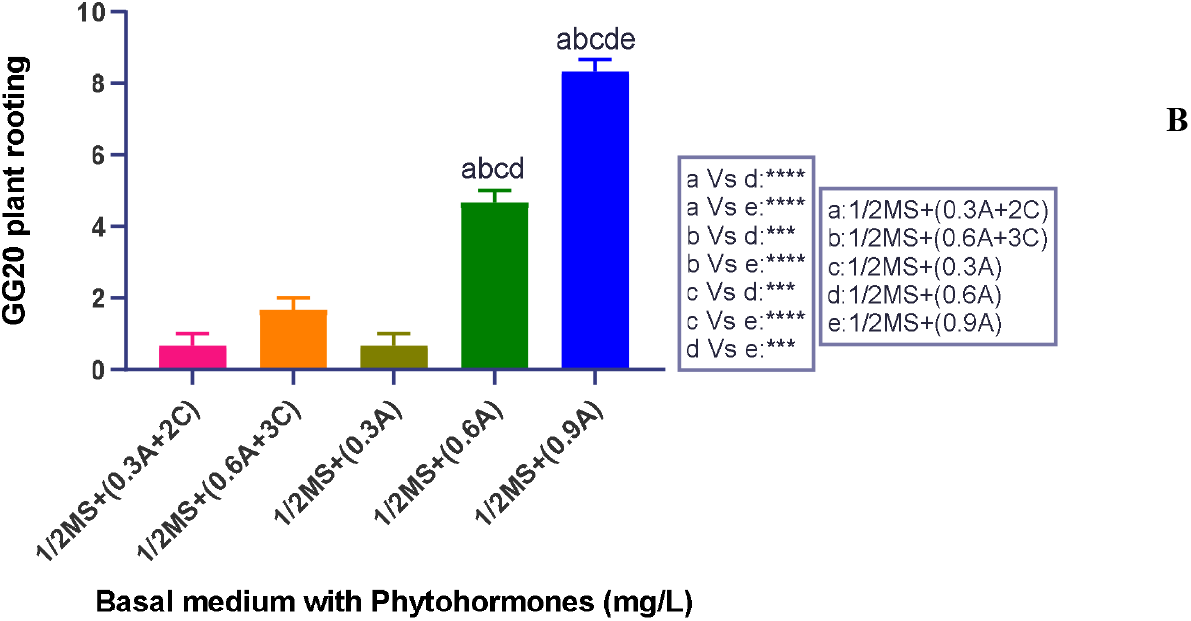
**A)** Effects of Basal media (MS) with (0.5Cmg/L) to (2Cmg/L) BAP on cotyledon node regeneration in GG20 (*Arachis hypogea*). MS: Murashige and Skoog medium; C: cytokinin phytohormone. Bars with different letters indicates significant differences (*p* ≤ 0.05), as analysed by one-way ANOVA and Tukey’s multiple range test. **B)** Effects of Basal medium (MS) with (0.1A+1A, 0.3A+2C, 0.6A+3C, 0.1A, 0.3A, 0.6Amg/L) on cotyledon node regenerated shoots in GG20 (*Arachis hypogea*). MS: Murashige and Skoog medium; A: auxin phytohormone (NAA); C: cytokinin phytohormone (BAP). Bars with different letters indicates significant differences (*p* ≤ 0.05), as analysed by one-way ANOVA and Tukey’s multiple range test.

**Figure 4.**
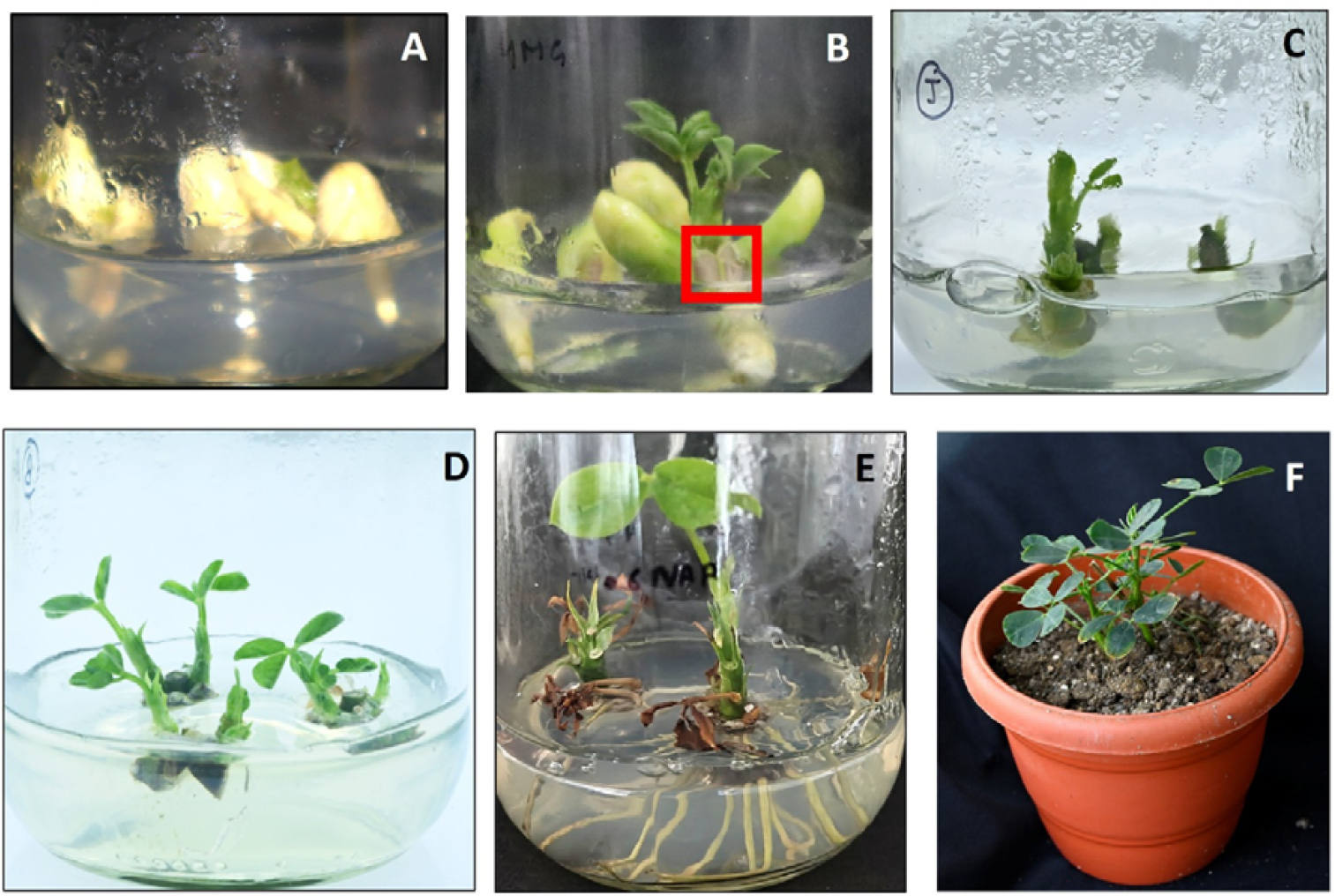
Tissue culture and hardening of *GJG9* peanut cultivar; **(A)** Sterilized *A. hypogaea GJG9* seed on MS media and seed germination starts after sterilization; **(B)** cotyledon node used for regeneration after seed germination; (Red mark represents that this portion excised by cutting and used as a (CNs) cotyledon node); **(C)** cotyledon nodes regenerate on MS+ 2mg/L BAP medium; **(D)** Multiple regenerated shoots were ready for rooting; **(E)** Regenerated shoots were used for rooting on MS+ 0.6 mg/L NAA medium; **(F)** Rooted plants transferred in pots for the hardening purpose. **(C)** Cotyledon nodes regenerate on MS+ 2mg/L BAP medium; (E) Regenerated shoots were used for rooting on MS+ 0.9 mg/L NAA medium; **(F)** Rooted plants transferred in pots for the hardening purpose.

**Figure 5.**
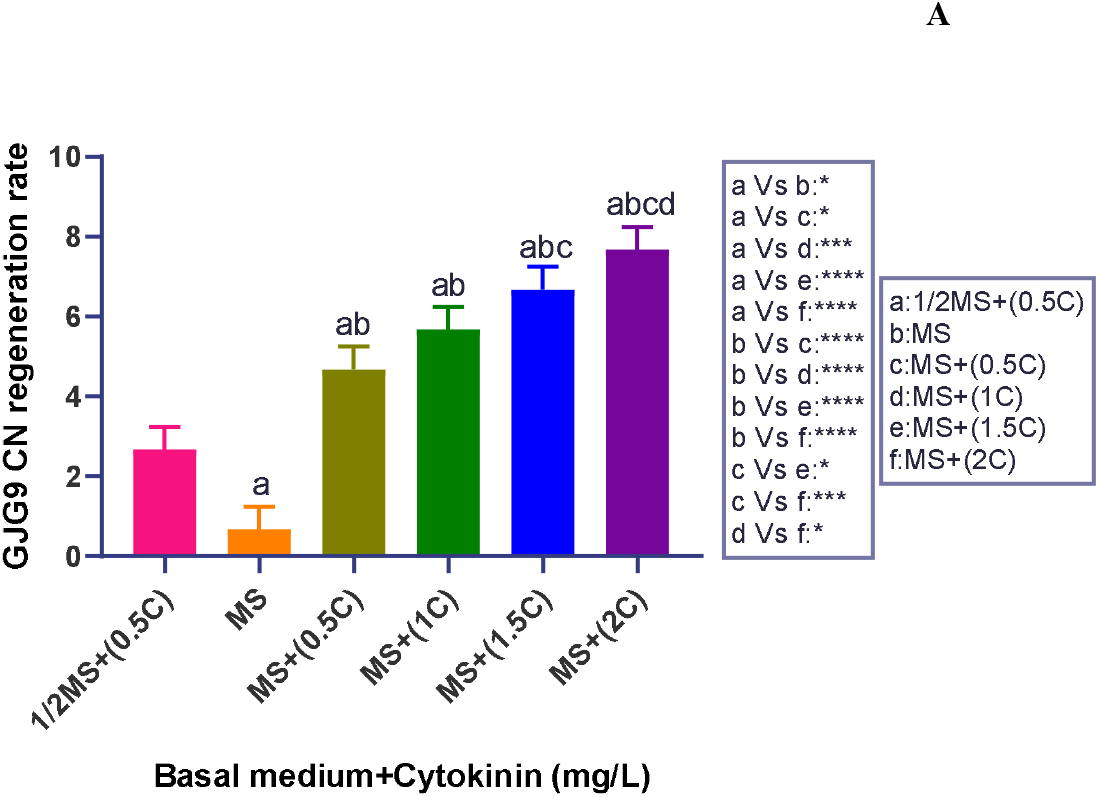
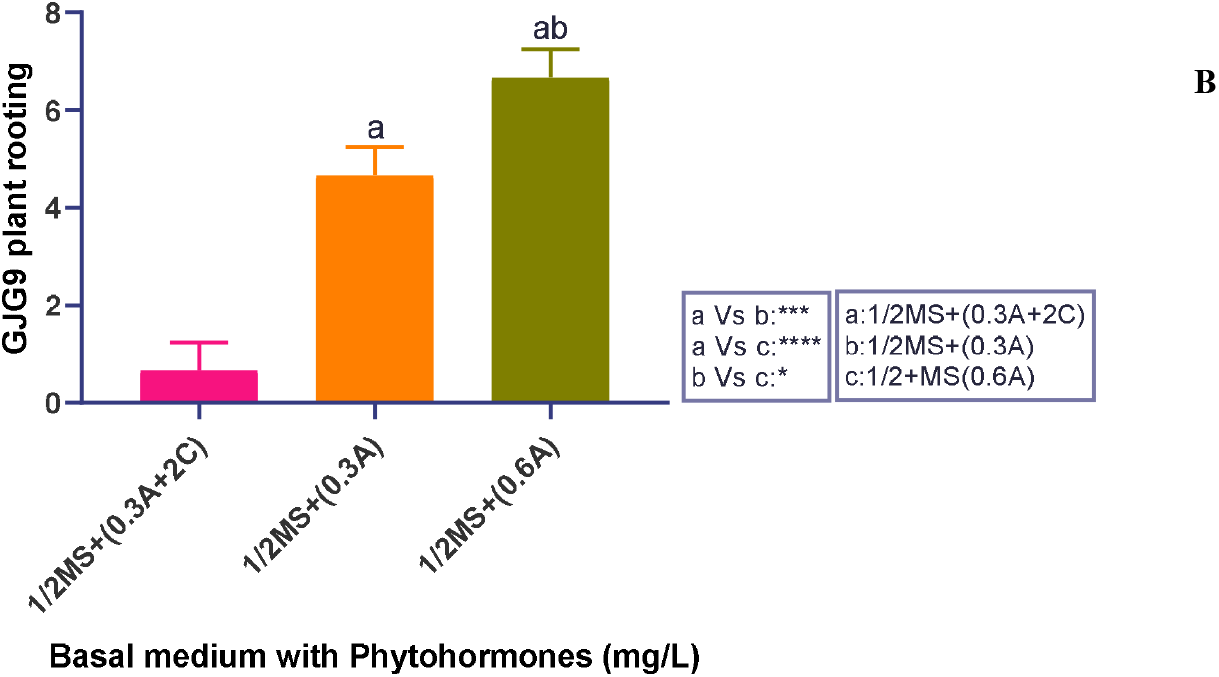
**A)** Effects of Basal media (MS) with (0.5Cmg/L) to (2Cmg/L) BAP on cotyledon node regeneration in GJG9 (*Arachis hypogea*). MS: Murashige and Skoog medium; C: cytokinin phytohormone. Bars with different letters indicates significant differences (*p* ≤ 0.05), as analysed by one-way ANOVA and Tukey’s multiple range test. **B)** Effects of Basal medium (MS) with (0.1A+1C, 0.3A+2C, 0.1A, 0.3A, 0.6A mg/L) on induced roots in GJG-9 shoots (Arachis hypogea). MS: Murashige and Skoog medium; A: auxin phytohormone (NAA); C: cytokinin phytohormone (BAP). Bars with different letters indicates significant differences (*p* ≤ 0.05), as analysed by one-way ANOVA and Tukey’s multiple range test.

**Figure 6.**
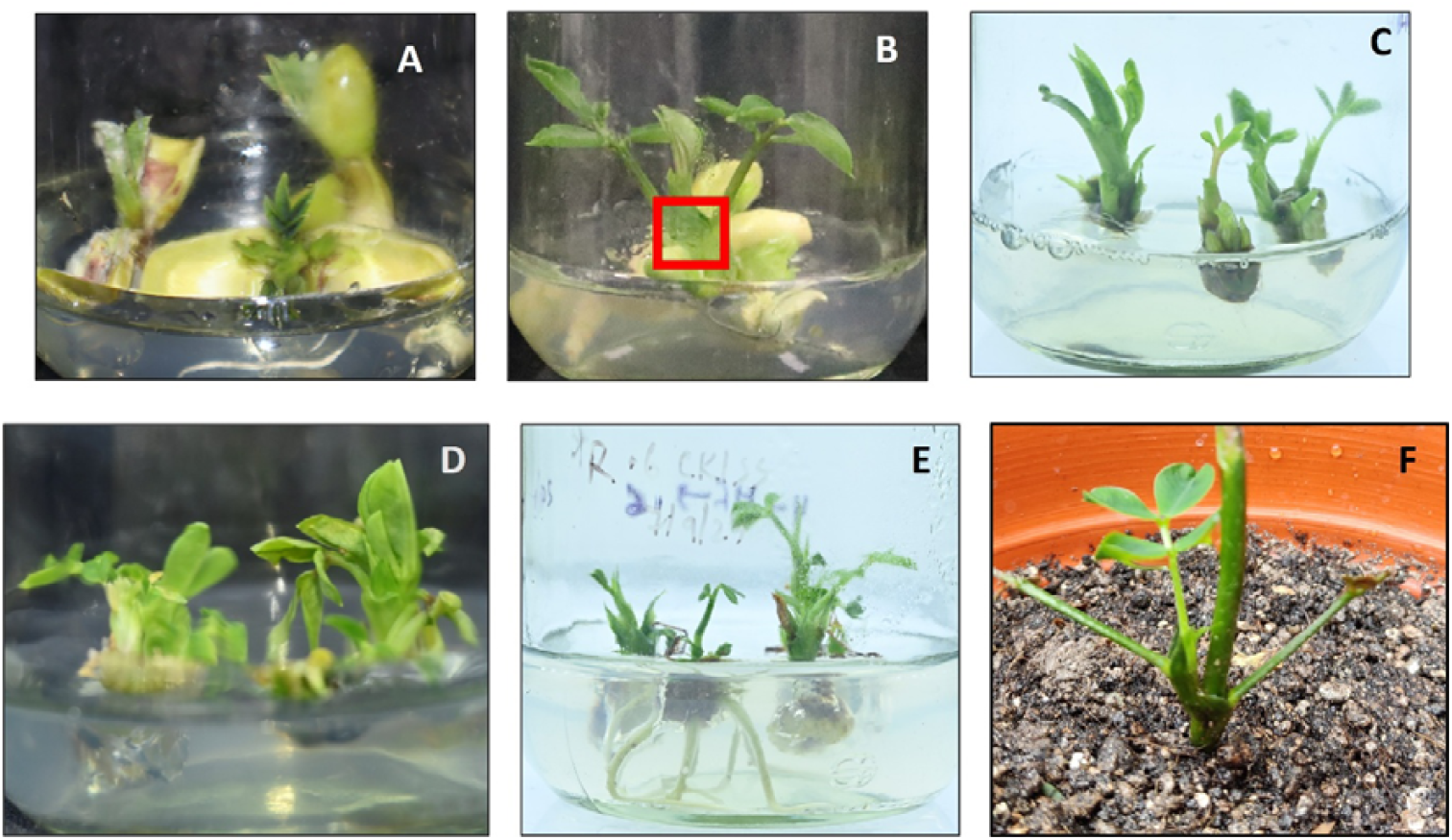
Tissue culture and hardening of TAG-37A peanut cultivar; **(A)** Sterilized *A. hypogaea TAG-37A* seed on MS media and seed germination starts after sterilization; **(B)** cotyledon node used for regeneration after seed germination; (Red mark represents that this portion excised by cutting and used as a (CNs) cotyledon node); **(C)** cotyledon nodes regenerate on MS+ 4mg/L BAP medium; **D)** Multiple regenerated shoots were ready for rooting; **(E)** Regenerated shoots were used for rooting on MS+ 0.9 mg/L NAA medium; **(F)** Rooted plants transferred in pots for the hardening purpose.

**Figure 7.**
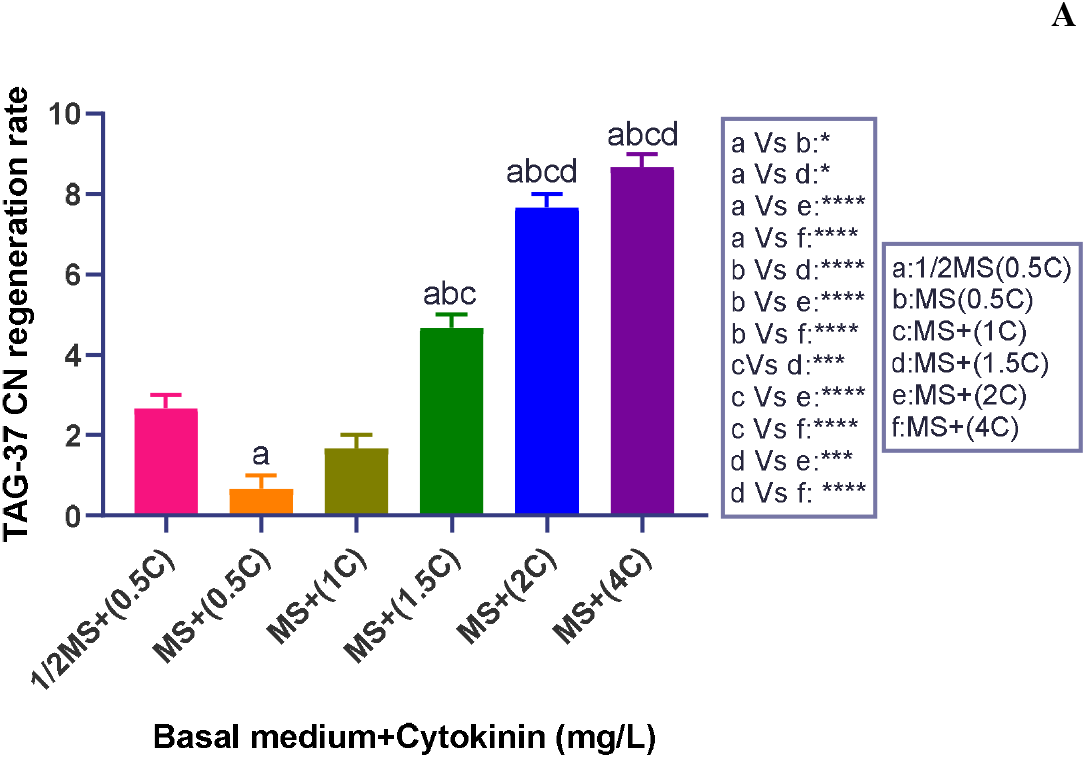
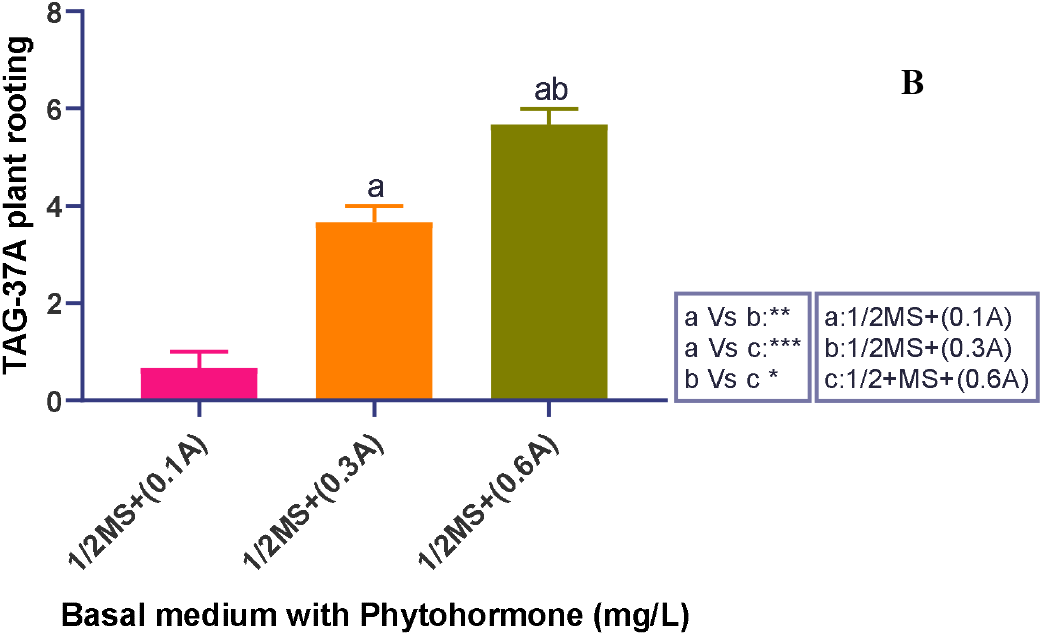
**A)** Effects of Basal media (MS) with (0.5Cmg/L) to (4Cmg/L) BAP on cotyledon node regeneration in TAG-37A (Arachis hypogea). MS: Murashige and Skoog medium; C: cytokinin phytohormone. Bars with different letters indicates significant differences (p ≤ 0.05), as analysed by one-way ANOVA and Tukey’s multiple range test. **B)** Effects of Basal medium (MS) with (0.1+1C, 0.3A+2C, 0.3A, 0.6Amg/L) on induced roots in TAG-37A shoots (Arachis hypogea). MS: Murashige and Skoog medium; A: auxin phytohormone (NAA); C: cytokinin phytohormone (BAP). Bars with different letters indicates significant differences (p ≤ 0.05), as analysed by one-way ANOVA and Tukey’s multiple range test.

**Table 2.** Optimization of cotyledon node regeneration rate of (*Arachis hypogea*); GG20 variant;

| S. No | MS media | Name of variant | Cotyledon node Transferred on Cytokinin (mg/L) | Cotyledon node Regeneration Efficiency (%) |
| --- | --- | --- | --- | --- |
| 1. | ½ Basal Media | GG-20 | 0 | 0 |
| 2. | ½ Basal Media | GG-20 | 0.5 | 33.33 |
| 3. | Full Basal Media | GG-20 | 0 | 11.11 |
| 4. | Full Basal Media | GG-20 | 0.5 | 55.55 |
| 5. | Full Basal Media | GG-20 | 1 | 66.66 |
| 6. | Full Basal Media | GG-20 | 1.5 | 77.77 |
| 7. | Full Basal Media | GG-20 | 2 | 88.88 |

**Table 3.** Optimization of rooting rate of shoots of (*Arachis hypogea*); GG20 variant.

| S. No | MS media | Name of variant | Auxin (mg/L) | Cytokinin (mg/L) | Rooting Efficiency (%) |
| --- | --- | --- | --- | --- | --- |
| 1. | ½ Basal Media | GG-20 | 0 | 0 | 0 |
| 2. | ½ Basal Media | GG-20 | 0.1 | 1 | 0 |
| 3. | ½ Basal Media | GG-20 | 0.3 | 2 | 11.11 |
| 4. | ½ Basal Media | GG-20 | 0.6 | 3 | 22.22 |
| 5. | ½ Basal Media | GG-20 | 0.1 | 0 | 0 |
| 6. | ½ Basal Media | GG-20 | 0.3 | 0 | 11.11 |
| 7. | ½ Basal Media | GG-20 | 0.6 | 0 | 55.55 |
| 8. | ½ Basal Media | GG-20 | 0.9 | 0 | 88.88 |

**Table 4.** Optimization of cotyledon node regeneration rate of (*Arachis hypogea*); *GJG9* variant.

| S. No | MS media | Name of variant | Cotyledon node Transferred on Cytokinin (mg/L) | Cotyledon node Regeneration Efficiency (%) |
| --- | --- | --- | --- | --- |
| 1. | ½ Basal Media | GJG-9 | 0 | 0 |
| 2. | ½ Basal Media | GJG-9 | 0.5 | 33.33 |
| 3. | Full Basal Media | GJG-9 | 0 | 11.11 |
| 4. | Full Basal Media | GJG-9 | 0.5 | 55.55 |
| 5. | Full Basal Media | GJG-9 | 1 | 66.66 |
| 6. | Full Basal Media | GJG-9 | 1.5 | 77.77 |
| 7. | Full Basal Media | GJG-9 | 2 | 88.88 |

**Table 5.** Optimization of rooting rate of shoots of (*Arachis hypogea*); *GJG9* variant.

| S. No | MS media | Name of variant | Auxin (mg/L) | Cytokinin (mg/L) | Rooting Efficiency (%) |
| --- | --- | --- | --- | --- | --- |
| 1. | ½ Basal Media | GJG-9 | 0 | 0 | 0 |
| 2. | ½ Basal Media | GJG-9 | 0.1 | 1 | 0 |
| 3. | ½ Basal Media | GJG-9 | 0.3 | 2 | 11.11 |
| 4. | ½ Basal Media | GJG-9 | 0.1 | 0 | 0 |
| 5. | ½ Basal Media | GJG-9 | 0.3 | 0 | 55.55 |
| 6. | ½ Basal Media | GJG-9 | 0.6 | 0 | 77.77 |

**Table 6.** Optimization of cotyledon node regeneration rate of (*Arachis hypogea*); *TAG-37A* variant.

| S. No | MS media | Name of variant | Cotyledon node Transferred on Cytokinin (mg/L) | Cotyledon node Regeneration Efficiency (%) |
| --- | --- | --- | --- | --- |
| 1. | ½ Basal Media | TAG-37A | 0 | 0 |
| 2. | ½ Basal Media | TAG-37A | 0.5 | 33.33 |
| 3. | Full Basal Media | TAG-37A | 0 | 0 |
| 4. | Full Basal Media | TAG-37A | 0.5 | 11.11 |
| 5. | Full Basal Media | TAG-37A | 1 | 11.11 |
| 6. | Full Basal Media | TAG-37A | 1.5 | 55.55 |
| 7. | Full Basal Media | TAG-37A | 2 | 77.77 |
| 8. | Full Basal Media | TAG-37A | 4 | 88.88 |

**Table 7.**
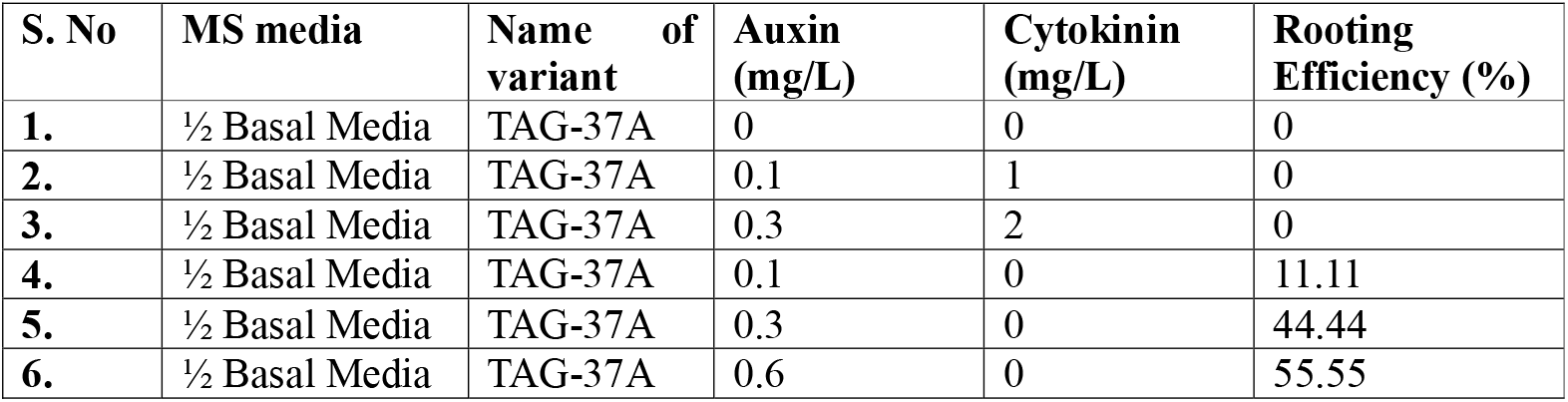
Optimization of rooting rate of shoots of (*Arachis hypogea*); TAG-37A variant.

| S. No | MS media | Name of variant | Auxin (mg/L) | Cytokinin (mg/L) | Rooting Efficiency (%) |
| --- | --- | --- | --- | --- | --- |
| 1. | ½ Basal Media | TAG-37A | 0 | 0 | 0 |
| 2. | ½ Basal Media | TAG-37A | 0.1 | 1 | 0 |
| 3. | ½ Basal Media | TAG-37A | 0.3 | 2 | 0 |
| 4. | ½ Basal Media | TAG-37A | 0.1 | 0 | 11.11 |
| 5. | ½ Basal Media | TAG-37A | 0.3 | 0 | 44.44 |
| 6. | ½ Basal Media | TAG-37A | 0.6 | 0 | 55.55 |

### Statistical analysis

The data were analyzed using analysis of variance (One way ANOVA) and Tukey’s multiple range test performed with the help of Prism software to determine the effects of cultivars for statistically significant. The *p-value* <0.05 was considered as the threshold for statistical significance between different data points.

### Agrobacterium-mediated transformation

*Agrobacterium tumefaciens* strain EHA105, harbouring the binary plant vector pGFPGUSPlus (addgene plasmid #64401) was used for transformation. This plasmid contained two reporter genes *EGFP* and *GUS* under control of CaMV35S promoter and nopaline synthase *(nos*) terminator and Hygromycin Phosphotransferase II (*HPTII*) as a selection marker gene.

*Agrobacterium* culture was prepared by growing a single colony in Luria Bertani broth medium and agar medium for 16-18 hours supplemented with rifampicin (25μg/ml) and kanamycin (50μg/ml) for selection.

### Media Composition

Liquid basal medium composition included MS major salts (Murashige and Skoog 1962), MS micronutrients and vitamins, 100 μM myo-inositol and acetosyringone, with pH adjusted to 5.7-5.8. The co-cultivation medium composition included MS major salts, MS micronutrients and vitamins, 100 μM myo-inositol, 3% sucrose, 0.8% agar and acetosyringone, with pH adjusted to 5.7-5.8.

### Culture method

For broth cultures, the bacterial suspension was adjusted to an OD□ □ □ of 0.6–0.8 and centrifuged at 5,000 rpm for 5 min to collect the cells. The pellet was re-suspended in 50 mL of liquid basal medium supplemented with acetosyringone (100 μM to 150 μM). Approximately 30–40 explants were immersed in the suspension and incubated for either 2 min or 15 min. Excess bacterial suspension was removed from the explants, followed by co-cultivation on co-cultivation medium supplemented with 100 μM acetosyringone for 2, 3, or 5 days under dark conditions.

After co-cultivation, explants were washed and transferred to plastic pots containing sterile Soilrite mix (Keltech Energies Ltd., Bangalore, India). Putative transformants were allowed to grow for 2–3 weeks under a 16:8 h light–dark photoperiod at 25 ± 2 °C before being transferred to larger pots (15–20 inch diameter) for further growth and maturation. Explants regenerated into complete plants in the absence of *Agrobacterium* infection were used as negative controls for subsequent experiments.

### Single colony method

A positive single colony from the culture plate was suspended in 50 ml of liquid MS basal medium supplemented with acetosyringone. This transformation procedure was carried out following the same protocol described for the culture method.

### Analyses of putative transgenic plants

Histochemical assays were performed to assess GUS activity in seeds (cotyledons) following incubation on co-cultivation medium and in leaves collected 1–2 weeks after transfer to plastic pots. GUS staining was carried out using X-Gluc (5-bromo-4-chloro-3-indolyl-β-D-glucuronide) as the substrate, following the protocol described by Rajput et al. (2024).

Total genomic DNA was isolated from fresh young leaves using CTAB method. Polymerase chain reaction (PCR) was performed with an initial denaturation at 95 °C for 5 min, followed by 35 cycles of denaturation at 95 °C for 35 s, annealing at 58 °C for 40 s, and extension at 72 °C for 45 s, with a final extension at 72 °C for 5 min. Amplification of the transgenes was carried out using gene-specific primers for the GUS gene (forward: 5′-ACC GCG ACG TCT GTC GAG-3′; reverse: 5′-CCA GTG ATA CAC ATG GGG ATC-3′) and the hygR gene (forward: 5′-TTC TTT GCC CTC GGA CGA GTG-3′; reverse: 5′-ACA GCG TCT CCG ACC TGA TG-3′). Plasmid DNA of pGFPGUSPlus was used as a positive control in all PCR analyses.

## Results

### Optimization of seed Germination

MS basal medium strength of significantly influenced in vitro seed germination efficiency in all three peanut cultivars (Fig. 1) (Table 1). In all genotypes, full-strength MS medium resulted in higher germination rate compared with half-strength MS medium. Among the cultivars tested, GJG-9 exhibited the highest germination efficiency (90%) on full-strength MS medium, followed by TAG-37A and GG-20, both recording 80% germination. On half-strength MS medium, germination efficiency ranged from 60% to 70%, with GJG-9 again showing comparatively higher response (Fig. 1) (Table 1). TAG-37A consistently showed improved germination on full-strength MS medium compared with half-strength medium, indicating a positive response to increased nutrient availability. The enhanced germination observed on full-strength MS medium resulted in vigorous and uniform seedlings, which were subsequently used for cotyledonary node (CN) explant preparation in regeneration experiments.

### Optimization of shooting and rooting media

The effect of MS basal medium and cytokinin phytohormone concentration on CNs mediated shoot regeneration were evaluated in three peanut cultivars: GG-20, GJG-9, and TAG-37A (Table. 2,4,6) (Fig. 3A, 5A, 7A). The CNs explant cultured on MS media without any phytohormone failed to regenerate shoots or showed only few responses, irrespective of medium strength, indicating that exogenous phytohormone, cytokinin was essential for shoot induction.

The effect of auxin (NAA) alone or in combination of cytokinin (BAP) supplementation on in vitro root induction from regenerated shoots was evaluated using half-strength MS basal medium in three peanut cultivars—GG-20, GJG-9, and TAG-37A. Rooting response was assessed as rooting efficiency (%) under different phytohormone combinations (Table. 3,5,7) (Fig. 3B, 5B, 7B).

### Response of GG-20

In CN no shoots formed on half-strength MS without cytokinin (BAP), however 0.5 mg/L BAP achieved 33.33% efficiency. Full-strength MS showed 11.11% basal regeneration, rising BAP concentration dependently to 55.55% (0.5 mg/L), 66.66% (1mg/L), 77.77% (1.5 mg/ L). The highest regeneration efficiency (88.88%) was achieved on full-strength MS medium supplemented with 2 mg/ L BAP (Table. 2) (Fig. 3A).

No root formation was observed on hormone-free medium or on media supplemented with low auxin NAA (0.1 mg/L) in combination with cytokinin in GG-20. Limited rooting response (11.11–22.22%) was recorded when auxin NAA was supplied together with cytokinin BAP at higher concentrations (Fig. 3B) (Table. 3). In contrast, media supplemented with auxin NAA alone resulted in a marked increase in rooting efficiency. Rooting frequency increased progressively with increasing auxin concentration, reaching 11.11% at 0.3 mg/L, 55.55% at 0.6 mg/L, and a maximum of 88.88% at 0.9 mg/L NAA. These results indicate that auxin alone was more effective than auxin–cytokinin combinations for root induction in GG-20 (Table. 3) (Fig. 3B).

### Response of GJG-9

A similar regeneration pattern was observed in GJG-9 as GG-20. Half-strength MS medium without BAP cytokinin failed to induce shoot regeneration from CNS, while the addition of 0.5 mg/L BAP resulted in 33.33% regeneration. On full-strength MS medium, regeneration efficiency increased in a cytokinin-dose-dependent manner, from 11.11% BAP (0 mg/L) to 55.55% (0.5 mg/L), 66.66% (1 mg/L), and 77.77% (1.5 mg/L). Maximum regeneration (88.88%) was recorded at 2 mg/L BAP (Table. 4) (Fig. 5 A), indicating high responsiveness of this genotype to moderate cytokinin levels.

No rooting was observed on hormone-free medium or in the presence of low auxin NAA combined with cytokinin BAP in GJG-9. A moderate rooting response (11.11%) was observed at higher auxin–cytokinin combinations. However, when auxin was supplied alone, rooting efficiency increased substantially. Rooting frequencies of 55.55% and 77.77% were recorded at 0.3 mg/L and 0.6 mg/L NAA auxin, respectively, indicating that GJG-9 exhibited high sensitivity to auxin-mediated root induction (Table. 5) (Fig. 4E, 5 B).

### Response of TAG-37A

In TAG-37A, CNs cultured on half basal MS medium without cytokinin showed no regeneration, while 0.5 mg/L BAP cytokinin resulted in 33.33% regeneration. Unlike GG-20 and GJG-9, TAG-37A exhibited a lower response at lower BAP cytokinin concentrations on full-strength MS medium, with only 11.11% regeneration at 0.5 mg/L and 22.22% at 1 mg/L. However, regeneration efficiency of CNs increased substantially at higher BAP concentrations, reaching 55.55% at 1.5 mg/L and 88.88% at 2 mg/L. The highest regeneration efficiency (99.99%) was achieved at 4 mg/L BAP, indicating that TAG-37A requires higher cytokinin levels for optimal shoot induction (Table. 6) (Fig. 7 A).

TAG-37A exhibited comparatively lower rooting efficiency than the other two cultivars. No rooting was observed on hormone-free medium or on media supplemented with auxin in combination with cytokinin. When auxin was supplied alone, rooting response improved gradually, with 11.11% rooting at 0.1 mg/L, 44.44% at 0.3 mg/L, and a maximum of 55.55% at 0.6 mg/L auxin NAA (Table. 7) (Fig. 7 B).

Across all three cultivars, full basal MS medium consistently supported higher CNs regeneration efficiency than half basal MS medium. CNs shoot regeneration was strongly dependent on cytokinin concentration, with a clear dose-dependent increase in regeneration frequency. However, the optimal cytokinin requirement varied among genotypes. GG-20 and GJG-9 showed maximum regeneration at 2 mg/L BAP, whereas TAG-37A required a higher concentration (4 mg/L) to achieve maximum regeneration.

However, in case of rooting in all three cultivars, half-strength MS medium supplemented with auxin alone was significantly more effective for root induction than media containing both auxin and cytokinin. Rooting efficiency increased in a dose-dependent manner with increasing auxin concentration, although the optimal auxin requirement varied among genotypes. GG-20 exhibited the highest rooting efficiency (88.88% at 0.9 mg/L NAA), followed by GJG-9 and TAG-37A.

### Development of plumular meristem transformation via *Agrobacterium* tumefaciens– mediated

Overnight soaked and surface sterilized seeds were needle-pricked at the plumular meristem junction near the embryo axis and subjected to *Agrobacterium* tumefaciens–mediated transformation (Fi**g** 8). Initial attempts using an optical density (OD)-based bacterial suspension resulted in excessive bacterial proliferation during co-cultivation, leading to tissue necrosis and reduced regeneration frequency. To overcome this limitation, a single-colony inoculation method was adopted, which significantly minimized bacterial overgrowth and improved explant survival. A co-cultivation period of 2–3 days was found to be optimal for efficient transformation, whereas prolonged co-cultivation adversely affected regeneration due to bacterial overgrowth. Following co-cultivation, seedlings were transferred to soilrite and maintained for 2–3 weeks, after which approximately 63.88% of the regenerated shoots were successfully established in soil. Using peanut cultivar GG-20, a total of eight putative transgenic plants were obtained across two independent experiments, with an average transformation efficiency of 7.69%, calculated as the percentage of T□ plants obtained relative to the total number of explants co-cultivated with A. tumefaciens (Table 8).

**Table 8:** Frequency of *A. tumefaciens*-mediated plumular meristem transformation in peanut cultivar GG-20.

| No. of batches | Explants Infected by *Agrobacterium* | No. of plants surviving in soilrite | No. of plants Analyzed by PCR | No. of positive plants | Transformation efficiency (%) |
| --- | --- | --- | --- | --- | --- |
| 5 | 225 | 128 | 92 | 8 | 8.70 |

**Figure 8.**
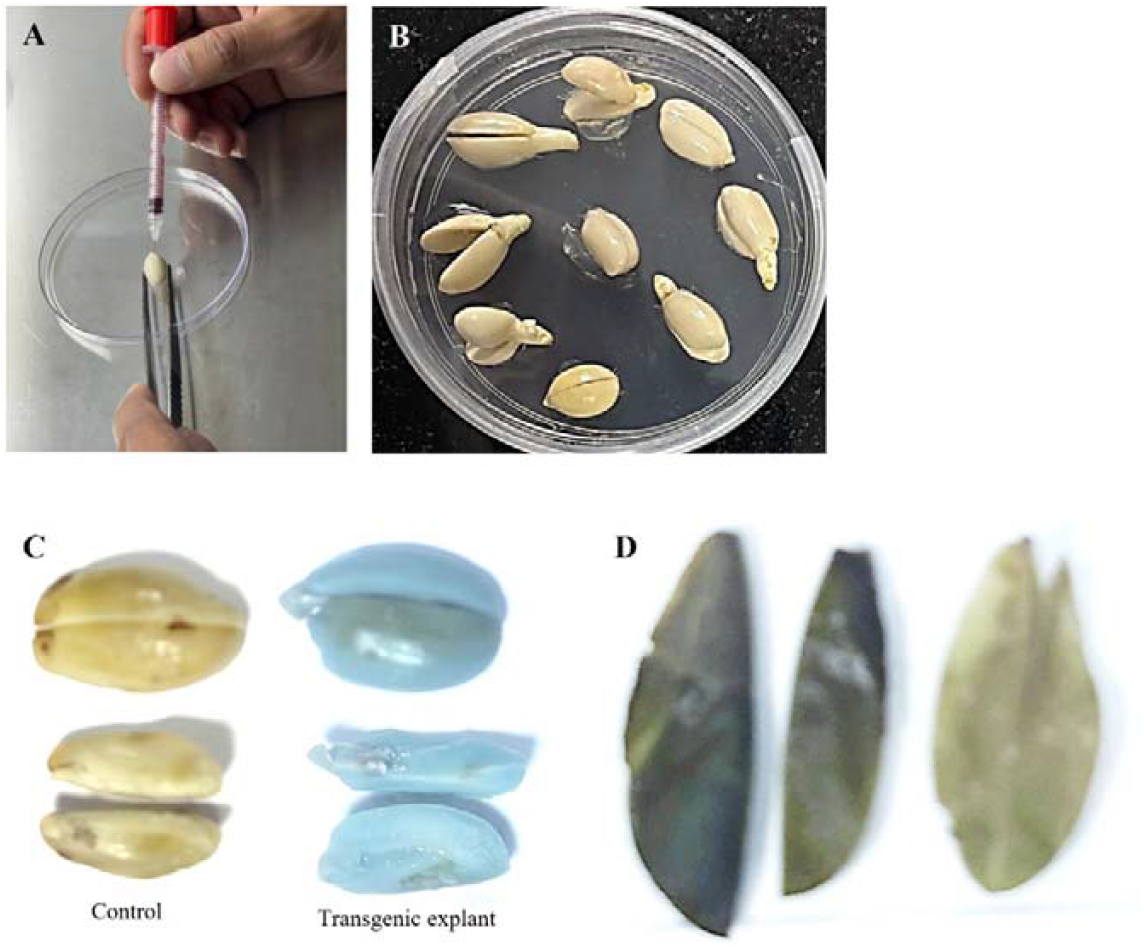
*Agrobacterium-*mediated transformation. **A**) GG20 seed pricking near the embryo; **B)** *Agrobacterium* transformed seeds in co-cultivation media. GUS histochemical assay. **C)** GUS staining of untransformed and transgenic seeds after co-cultivation; **D)** GUS staining of transgenic (left two leaves) and control leaf (towards the right), 1-2 weeks after transfer into soil.

### GUS expression and PCR analysis of selected T0 plants

Histochemical GUS staining of the explants at various developmental stages to screen putative transgenic plants showed blue colouration of the transformed shoots whereas no GUS activity was observed in untransformed control (Fig. 8C, D). GUS analysis was done in embryo after co-cultivation and leaves after 2 and 4 weeks of growth under soilrite.

Ninety two independent primary peanut transformants were established using *Agrobacterium*-mediated plumular meristem transformation method (Fig. 9). PCR based molecular screening was carried out initially to detect the presence of transgene in T_0_ events. The 8 established T_0_ progeny lines were confirmed through PCR analysis for the presence of both *HPTII* of 468 bp (Fig. 10) and *GUS* of 516 bp (Fig. 11) genes.

**Figure 9.**
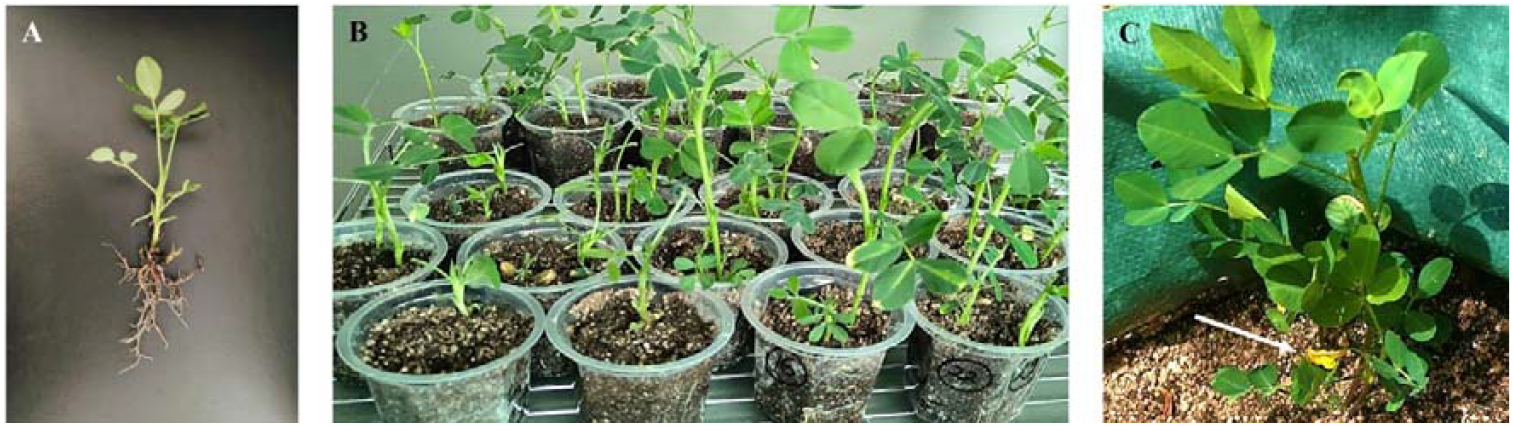
Different stages of hardening and transplantation. **A)** 1-2 weeks seedling after co-cultivation with visible secondary root formation; **B)** Plants growing in culture room after 3 weeks of soil transfer; **C)** Established plant with flowers at green house after 8 weeks of soil transfer (arrow points at flowering).

**Figure 10.**
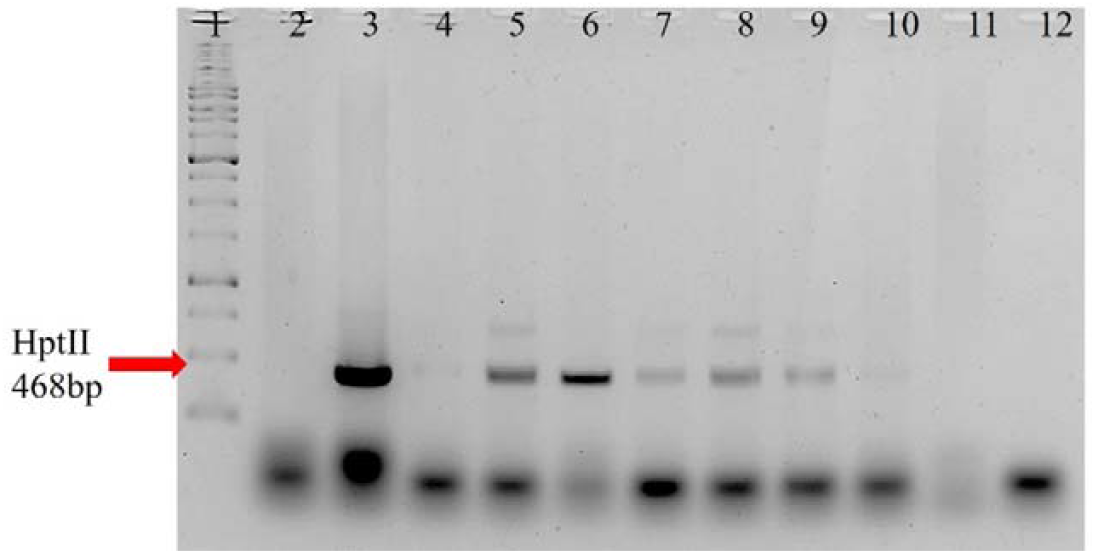
PCR analysis of established putative transformed plants for the presence of *hptII* gene. H) Amplification of 468 bp *hptII* specific fragment in T□plants; lanes 4-12. Lanes 1-3, DNA ladder, negative (Wt) and positive controls respectively.

**Figure 11.**
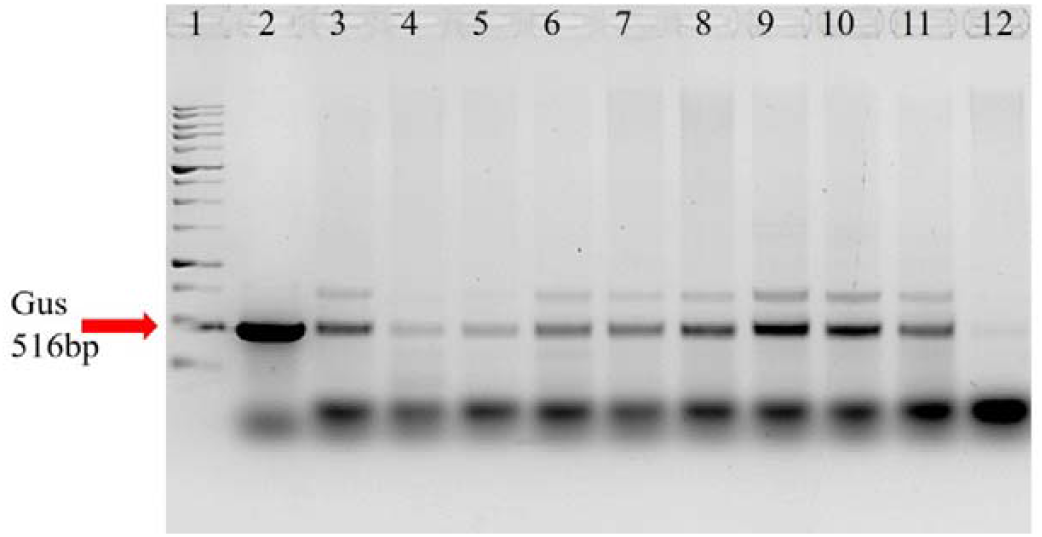
PCR analysis of established putative transformed plants for the presence of GUS gene. I) Amplification of 516 bp specific fragment in T□plants; lanes 3-11. Lanes 1-2 and 12, DNA ladder, positive and negative (Wt) controls respectively.

## Discussion

This study establishes a reproducible in vitro regeneration system using cotyledonary node (CN) explants from three peanut cultivars, revealing pronounced genotype-dependent responses in shoot organogenesis and root induction. CN explants derived from young seedlings exhibited high morphogenic competence, consistent with earlier reports indicating that nodal tissues retain pre-existing meristematic cells capable of direct organogenesis (Cheng et al., 1992; Radhakrishnan et al., 2000; Anuradha et al., 2006; Iqbal et al., 2012; Hsieh et al., 2017).

Shoot regeneration efficiency increased with increasing BAP (cytokinin) concentration across all GG-20, GJG9 and TAG-37A peanut cultivars, but optimal hormonal levels varied. GG-20 and GJG-9 exhibited maximal cotyledonary node regeneration (88.9%) at moderate cytokinin levels (2 mg/L), whereas TAG-37A required a substantially higher concentration (4 mg/L) to achieve near-complete shoot induction (∼89-100%). Such genotype-specific cytokinin responsiveness has been widely reported in peanut and other legumes and is commonly attributed to variation in endogenous hormone balance, cytokinin perception, and downstream signaling pathways (Arun et al., 2014; Hsieh et al., 2016). The utilization of authentic CN explants facilitates rapid and genotype-independent shoot regeneration (Sanyal et al. 2003; Hsieh et al., 2016), while CNs with exposed meristematic cells serve as valuable resources for Agrobacterium-mediated or biolistic transformation (Sanyal et al. 2003; Hsieh et al., 2016).

Overall, shoot induction rates ranging from 43–97% have been reported when CNs explants were used with optimized regeneration protocols across different legume species. In our CNs based protocol, the shoot induction frequency ranged from 88 to 99% in all cultivars, which is substantially higher than the 25% shoot induction reported previously in a peanut cotyledon regeneration system (Anuradha et al., 2006; Burns et al., 2012). While auxins are frequently applied to stimulate rooting, high concentrations can suppress shoot development and cause leaf abscission. Consequently, using reduced auxin levels is necessary to maintain a balance between root formation and shoot growth, emphasizing the importance of optimizing hormone concentrations for efficient regeneration (Anuradha et al., 2006; Hsieh et al., 2016). Previous studies have reported that NAA concentrations ranging from 0 to 5.37 μM are optimal for root induction in peanut (Anuradha et al., 2006; Burns et al., 2012; Hsieh et al., 2016). The present study among the cultivars evaluated, GG-20 showed the highest rooting efficiency, with nearly 90% rooting at higher auxin (0.6-0.9 mg/L NAA) concentrations, followed by GJG-9 and TAG-37A. The relatively lower rooting response observed in TAG-37A indicates possible genotype-specific differences in auxin perception or transport, which may affect adventitious root initiation and development.

Despite significant advances in genetic engineering, the development and exploitation of transgenic peanut lines remain limited compared to other major oilseed crops such as soybean and canola, largely due to recalcitrant transformation responses and strong genotype dependence. The present study also demonstrates the successful development of a plumular meristem–mediated *Agrobacterium* transformation protocol in peanut cultivar GG-20. A co-cultivation period of 2–3 days was found to be optimal for achieving efficient transformation, as longer durations resulted in excessive bacterial overgrowth and reduced regeneration frequency. Similar observations have been reported in earlier studies, where prolonged co-cultivation negatively affected explant viability and shoot regeneration due to bacterial toxicity. The high establishment rate of regenerated shoots (approximately 90%) indicates that the protocol is suitable for efficient recovery of transformed plants under soilrite conditions. The average transformation efficiency of 7.69% obtained in the present study is comparable to, or higher than, previously reported efficiencies in peanut transformation using meristem-based approaches.

In summary, this study delivers genotype-optimized regeneration and *in-planta* transformation protocols that overcome peanut’s recalcitrance, enabling precise genetic improvements. These tools will accelerate functional genomics, stress-resilient cultivars, and enhanced oil/nutritional quality to meet global demands. Future applications in CRISPR editing and multi-trait stacking promise transformative impacts on peanut productivity and food security.

## Author contributions

CK: Designed study, performed tissue culture experiments, writing-original draft, PR: Developed the *in planta* transformation protocol, genotyping, helped with manuscript draft; H G helped in tissue culture, *in-planta* transformation protocol experiments and genotyping, MP, MS, KHN and LP: Contributed to tissue culture optimization, MJ: Provided suggestions for tissue culture protocols and design of study, SS: Conceptualization, Funding acquisition, Supervision, Writing and Editing, BP: Conceptualization, Funding acquisition, Supervision, Writing and Editing.

## Funding

The author(s) declare that financial support was received for the research, authorship, and/or publication of this article. This work is supported by Gujarat State Biotechnology Mission, Govt. of Gujarat’s research grant to BP and SS under the Research Support Scheme 2022-23, project # GSBTM/JD (R&D)/626/22-23/00018354.

## Acknowledgments

BP acknowledges Ahmedabad University for the start-up funds. SS acknowledges DBT for the Re-entry fellowship grant, SERB and the Indian Institute of Technology Gandhinagar for start-up grant. CK and PR acknowledge RA fellowships from GSBTM. We also thank and acknowledge: Dr. R Madariya, Dr. N.D. Dholariya and Dr. K.L. Dobariya from Junagadh Agricultural University for the GG-20 seeds; Alpgiri seeds for GJG-9 (drought-tolerant Spanish bunch), and TAG-37A, and Dr. Sangram Lenka from Gujarat Biotechnology University for advise on tissue culture. We also thank to Dhruvisha Bhanderi from Atmiya University and Pouras Desai from University of Toronto for technical help in the *in-planta* protocol optimization

## Notes

### Competing Interest Statement

The authors have declared no competing interest.

